# *Streptococcus pyogenes* infects human endometrium by limiting its immune response

**DOI:** 10.1101/713875

**Authors:** Antonin Weckel, Thomas Guilbert, Clara Lambert, Céline Plainvert, François Goffinet, Claire Poyart, Céline Méhats, Agnès Fouet

## Abstract

Group A *Streptococcus* (GAS), a Gram-positive human-specific pathogen yields 517,000 deaths annually worldwide, including 163,000 due to invasive infections and among them puerperal fever. GAS is their most feared etiologic agent. Puerperal fever still accounts for more than 75,000 maternal deaths annually and before the introduction of efficient prophylactic measures 10% childbirths were followed by the mother’s death. Yet little is known regarding GAS invasive infection establishment or GAS efficiency in causing postpartum infection. To characterize its early steps, we set up coordinated analyses of *ex vivo* infection of the human decidua, the puerperal fever portal of entry. We analyzed GAS behavior and the immune response triggered. We demonstrate that GAS (i) benefits from tissue secreted products to multiply; (ii) invades the tissue and leads to the death of half the cells within two hours *via* SpeB protease and Streptolysin O activities, respectively; (iii) impairs the tissue immune response. Immune impairment occurs both at the RNA level, with the induction of only a restricted immediate innate immune response, and at the protein level, in a SLO- and SpeB-dependent manner. Our study indicates that GAS efficient decidua invasion and immune response restraint favor its propensity to develop rapid invasive infections in a gynecological-obstetrical context.

## Introduction

Puerperal fever currently accounts for > 75,000 maternal deaths annually. *Streptococcus pyogenes*, also known as Group A *Streptococcus* (GAS), a Gram-positive human-specific pathogen, is its most feared etiologic agent (1). Between 2010-2016, GAS was responsible for 56 and 109 cases/100 000 persons per year of postpartum infections in the USA and the UK, respectively (2,3). In France, 65% and 55% of GAS invasive infections occurred in the gyneco-obstetrical sphere for women between 18- and 40-years of age, in 2017 and 2018, respectively. In addition to puerperal fever, GAS causes a wide range of infections, from non-invasive infections, such as pharyngitis and impetigo, to invasive infections, such as necrotizing fasciitis, bacteremia and streptococcic toxic shock syndrome, responsible altogether for 163,000 deaths annually (4). Finally, post-streptococcal sequelae, such as glomerulonephritis and rheumatic arthritis, are auto-immune diseases leading to 354,000 deaths annually. Virulence factors contributing to invasive diseases, including the extracellular cysteine protease SpeB and cytotoxin Streptolysin O (SLO), have been described *in vitro* and confirmed *in vivo* in animal models of infection (5–7). SpeB cleaves many host molecules, degrading, for example, fibronectin, fibrinogen, plasminogen and vitronectin, or activating, for example, matrix metalloproteases and IL-1β (for review (8)). SLO, in addition to being a pore forming toxin, facilitates the entry of the NAD glycohydrolase into eukaryotic cells and together these toxins contribute to cell lysis and to GAS pathogenesis (9,10).

In contrast to commensals or other pathogens, GAS does not colonize the vagina in the steady state (11). Yet before the introduction of efficient hygienic prophylactic measures, up to 10% post-partum women died of puerperal fever in lying-in hospitals. Furthermore, half of the cases of GAS-related puerperal fever occur within the first two days postpartum, with very rapid progression to severe sepsis (12). Hence, we hypothesized that GAS efficiently and promptly infects the decidua. The decidua is the mucosal uterine-lining during pregnancy and after childbirth. This tissue is made up of endometrial decidualized stromal cells, not epithelial cells, and 40% of resident leucocytes (13). Our team already showed that GAS readily adheres to primary human decidual stromal cells (14). However, to our knowledge, the early steps of human decidua infection by GAS or even, on a broader outlook, of GAS tissue invasion have never been characterized *ex vivo* on a human tissue.

The current model describing the invasive infections early steps, supported by studies on an infection model of 3D organotypic human skin, suggests adhesion to host tissue, growth, invasion of the tissue within 24 h pi (15). These are followed by resistance to immune responses, tissue destruction and systemic GAS infection (16). The observation of necrotizing fasciitis biopsy support the *in vivo* relevance of these results (15). However, questions remain regarding the events occurring during the first hours of infection of the tissues, particularly whenever GAS is in contact with stromal tissues such as the decidua. We aimed at characterizing these events, at identifying causes of the apparent efficiency of GAS to rapidly elicit puerperal fever and at unraveling mechanisms involved.

To that effect, we resorted to infect *ex vivo* human decidua and to characterize different aspects of the infection *via* multiple approaches. Since the immunological status of the decidua at the time of birth is particular (17), we infected *ex vivo* the upper portion of the decidua that is sloughed off with the placenta (18). The experiments were carried out on a timescale ranging from 15 min to several hours and we analyzed the different steps of the process relevant to invasive infections. Also, GAS is genetically diverse, up to 15% of the genome being composed of genetic mobile elements that furthermore carry virulence factors (19). Strain genotyping relies on the sequence of the 5’-end of the *emm* gene and more than 200 *emm*-types exist (20). *emm28* strains are associated with obstetrical and gynecological infections and the *emm28*-type is among the three most prevalent *emm*-types involved in invasive infections (21–23). We therefore infected the decidua with a wild-type (WT) *emm28* clinical strain isolated from a puerperal fever and its GFP and mutant derivatives (24).

We show that GAS benefits from tissue secreted product to multiply at the decidual surface, that it invades the tissue in the first 8 hours of infection and that it kills half of the cells within two hours. The role of two major virulence factors, SpeB and SLO, on these phenotypes is documented; SpeB is involved in the tissue invasion and SLO in the cytotoxicity. The tissue immune response is also characterized at transcriptional and protein accumulation levels, with SpeB and SLO modifying cytokine accumulation.

## Results

### GAS grows on human decidua during early steps of infection

Human decidual explants were infected with a wild-type isolate derivative expressing GFP (GFP-WT) (Figure 1). We used two complementary approaches to follow tissue colonization and invasion: either the tissue was infected under static conditions (sc) and fixed at different time points or, for live microscopy, under flow conditions (fc). GAS efficiently adheres to the tissue surface that contains a fibronectin layer covering the decidual cells. These cells are embedded in an extracellular matrix containing type IV collagen (Figure 1A). Furthermore, at 24 h pi sc, GAS formed microcolonies of up to 16 µm thick (Figure 1B) indicating the successful tissue surface colonization by GAS. We quantified GAS multiplication at the tissue surface during the first hours pi fc. In contact with 5 out of 6 tissues (Table S1), GAS multiplies until it covers the whole tissue surface (Figure 1C, 1D, Supp video 1). We measured the growth parameters of initially scattered bacteria. GAS grows exponentially at the tissue surface with the colonized surface doubling every 82 minutes (Figure 1E, 1F, Supp video 2). We also quantified the increase in bacterial thickness (Figure 1G, 1H). Its mean value doubled in 4 hours reaching a mean value of 4.2 µm (Figure 1H). For any given subject, the increase in thickness was locally heterogeneous with, after 6 h, hot-spots of more than 10 µm thick (Figure 1G). Yet, the overall thickness increase was alike on all subjects considered (Figure 1G, 1H). Hence, GAS grows in all three dimensions, with an apparent doubling time slightly below 80 min, accounting for its robust capacity to infect the human decidua.

**Figure 1.**
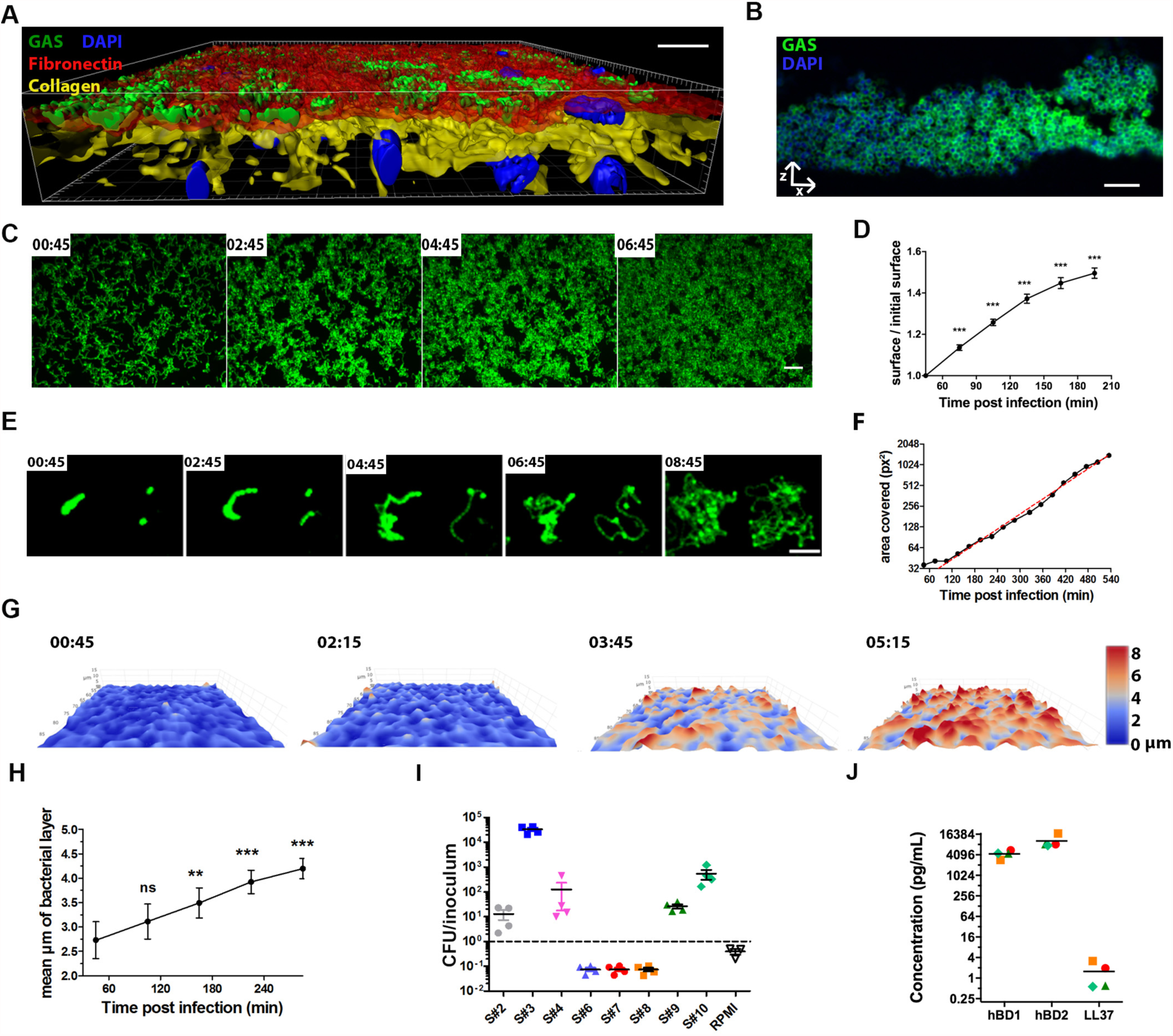
GAS adheres to the decidua and grows on it using tissue secreted products. **A**, Imaris 3D representation of a tissue infected 16 h sc and imaged *en face*. Fibronectin, red; type IV collagen, yellow; GFP-GAS, green; DAPI, blue. Scale bar: 20 µm. Magnification: 40X. **B**, Immuno-histofluorescence of a paraffin embedded tissue transversal slice (24 h pi, sc). Anti-GAS, green; DAPI, blue. Scale bar: 10 µm. Magnification: 100 X. **C** and **E**, z-max intensity projections of GFP signals from live fc *en face* acquisition images of GFP-WT at the tissue surface, at indicated time points, h:min. Scale bars: (**C**) 10 µm, (**E**) 20 µm. Magnification: 25 X. **D**, overtime ratio of the area covered by WT GAS GFP signal area in tissues **C** and from four other subjects compared to initial covered surface. Mean values of 3 to 11 fields for each of the 5 tissues are plotted. Two-Way ANOVA to the first time point. **F**, Quantification of the surface covered by GFP GAS from image (**E**); red dotted line, exponential curve fitted starting at 105 min. R^2^=0.9948. **G**, 3D surface heat-map of the bacterial layer thickness at different time-points (live imaging, fc) h:min. x, y and z are scaled, color code in µm. Image size: 303 x 190 µm. **H**, Quantification of the bacterial layer mean thickness, mean values of 2 to 13 fields for each of the 5 tissues are plotted. Two-Way ANOVA to the first time point. **I**, Multiplication factor after GAS incubation for 8 h in tissue-conditioned medium or RPMI. **J**, Basal levels of accumulation of the indicated antimicrobial peptide in the supernatant of non-infected decidual tissues 8 h after addition of fresh RPMI. Symbols same as in Figure 1I. For all panels, error bars are standard-error of the mean; values: ***, p<0.001, **, p<0.005.

To analyze if diffusible host factors promoted GAS growth, we compared GAS growth in the infection medium, RPMI, and in tissue-conditioned supernatants (Figure 1I). Whereas GAS does not multiply in RPMI, it multiplied in conditioned supernatants from 5 subjects and died in the other three (Table S1, Figure 1I). This suggests that, and non-exclusively, some tissues secreted nutrients and some bactericidal molecules. The antimicrobial peptides (AMP) human defensins (hBD) 1 and 2 and LL-37 were previously shown to be present in decidua during pregnancy (25). To define whether their amount could explain GAS growth limitation, we assayed the concentration of these AMPs in conditioned supernatants (Figure 1J). The hBD1 and hBD2 amounts were similar for all uninfected supernatants and that of LL-37 not significantly different, thus AMPs concentration do not seem predictive of growth possibility in the supernatants (Figure 1I).

### GAS invades the human decidua ex vivo

We then characterized GAS penetration of the decidua. While GAS is non-motile, bacteria were readily 4-5 micrometers below the tissue surface at 16 h pi sc (Figure 2A, S1A). To analyze the propensity of such events, we quantified invasion events through automated image analysis. As soon as 4 h pi sc, there are an average of 600 invasion events per mm^2^ (Figure 2B); this corresponds to approximately two invading chains per 1000 surface chains. Furthermore, there are three times more intra-tissular bacteria at 8 h than at 4 h pi (Figure 2C). We hypothesized that GAS invasion is partly due to tissue degradation, especially fibronectin degradation, and that SpeB, GAS major protease which degrades fibronectin (26), is involved in tissue penetration. To test this hypothesis, we infected decidua with either the WT or a ΔSpeB strain. At 4 h pi the ΔSpeB strain invaded the tissue two-fold less efficiently than the WT strain demonstrating that SpeB contributes to tissue invasion (Figure 2C). This suggests a possible involvement of matrix protein degradation in tissue invasion. Another potential source of bacterial invasion is internalization into host cells. We observed phagocytosis of GAS by immune cells 15 min pi fc and bacteria were found inside immune cells 6 µm below the tissue surface up to at least 4 h pi (Figure 2D, 2E, S1B), indicating that GAS may invade the tissue as intra-cellular bacteria. Our results indicate that GAS can rapidly and actively invade the decidua by an SpeB-dependent mechanism and may exploit survival in decidual immune cells.

**Figure 2.**
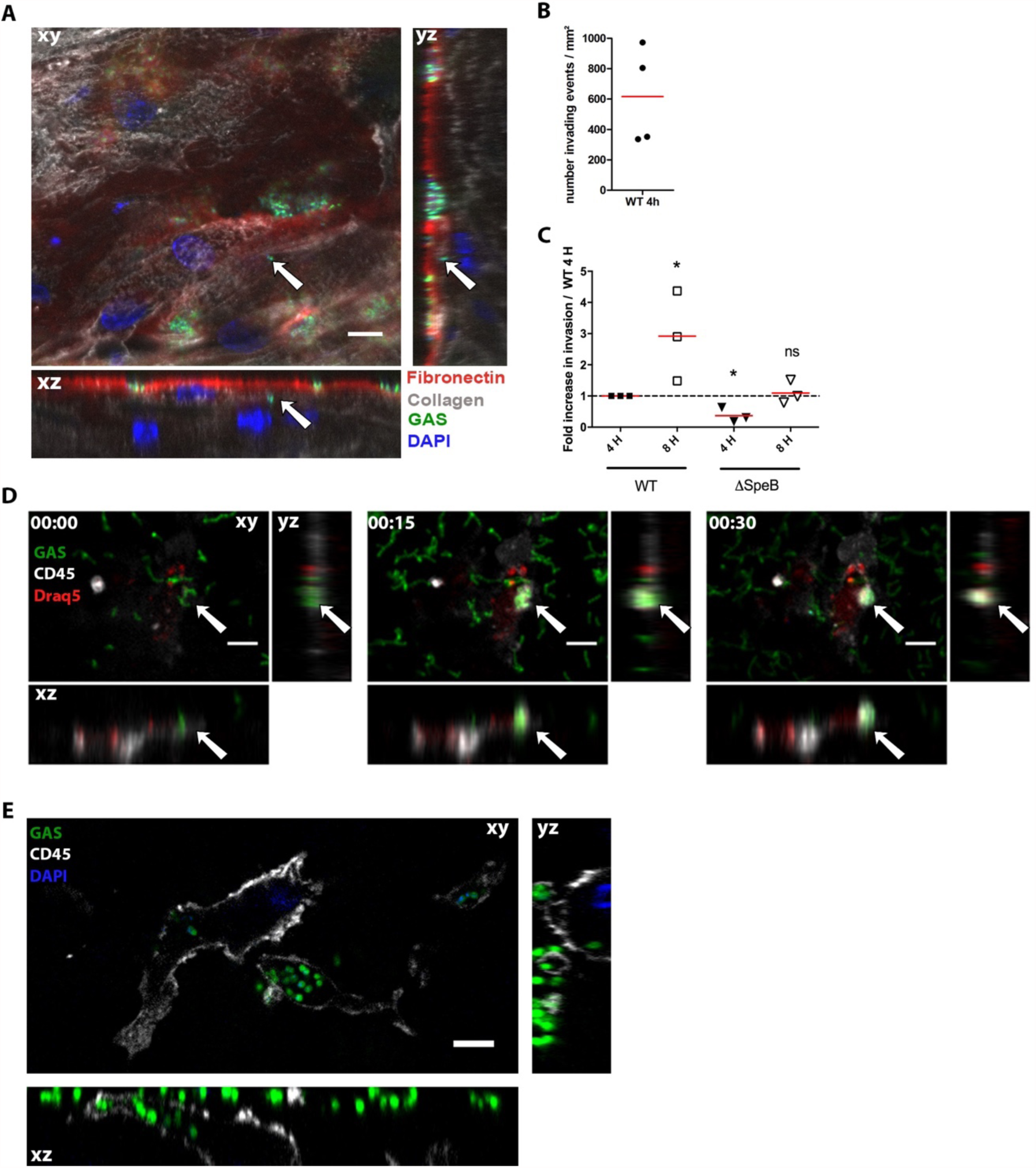
Protease activity and persistence in decidual immune cells permit GAS tissue invasion. **A**, Orthogonal view of a tissue 16 h pi sc. Arrows indicate an invading particle. Fibronectin, red; type IV collagen, grey; GAS, green; DAPI, blue. Scale bar: 10 µm. Magnification: 40 X. **B**, Mean number of invading events per mm^2^ at 4 h pi sc with the GFP-WT strain calculated using high throughput image analysis. Four subjects are shown. **C**, Ratio of GFP-WT and GFP-ΔSpeB invading events 4 and 8 h pi sc. One-sample t-test to the WT at 4 h for WT 8 h and ΔSpeB 4 h and to ΔSpeB 4 h for ΔSpeB 8 h. N=3. *, p<0.05. **D**, Time-lapse (fc) of an immune cell phagocytosing GAS *in situ*, with orthogonal view. Anti-CD45 (immune cell), grey; GAS, green; Draq5 (DNA), red. White arrows indicate the localization of the phagocytosis event. Scale bar: 10 µm. Magnification: 25 X**. E**, Orthogonal views of GFP-WT bacteria inside the tissue and within an immune cell; image taken 6 µm under the tissue surface 3 h pi fc. Anti-CD45, grey; bacteria, green; DAPI, blue. Scale bar: 5 µm. Magnification: 100 X.

### GAS induces cell death

GAS is known to induce cytotoxicity (27). We thus analyzed the consequences of *ex vivo* GAS infection of the human decidua on the physical state of cells. As soon as 4 h pi sc, we observed, as indicated by CD45 membrane protein visualization and DAPI staining, blebbing and dying immune cells (Figure 3A, Supp video 3) and at 16 h pi sc, by TUNEL assay, nuclei with altered morphology and DNA damages (Figure 3B). All cell-types, immune (CD45^+^) and non-immune (CD45^−^), were damaged (TUNEL^+^) (Figure 3C). We quantified the cytotoxicity induced by the infection at 2 and 8 h pi (Figure 3D, 3E). Twenty-seven % and 57% of cells were TUNEL positive at 2 h and 8 h pi, respectively, compared to 7% and 11% without infection (Figure 3E). To determine if cell death was induced by a GAS secreted factor, we measured cytotoxicity when bacteria were physically isolated from the tissue via a Transwell® insert (Figure 3E). In these conditions, GAS was still able to induce 21% and 37% of cell death at 2 h and 8 h pi, respectively, which demonstrates that GAS secreted factors induce cell death. Yet these values are lower than when bacteria are in contact with the tissues suggesting that direct contact or proximity is involved in the cytotoxicity efficiency. SLO (28,29) being a pore-forming toxin described to induce cell death, we tested to which extent this protein was involved in GAS cytotoxicity on the whole tissue at 2 and 4 h pi, infecting both primary decidual stromal cells and the decidua (Figure 3F, 3G). On primary decidual cells at 4 h pi, the WT and isogenic revertant (BTSLO) strains induced 60 % and 54% cytotoxicity, the isogenic SLO deleted mutant strain (ΔSLO) only 14%; in the non-infected condition there was only 7% cytotoxicity (Figure 3F). Hence, SLO is involved in the GAS cytotoxic effect. Also, on the decidual tissue, at 2 and 4 h pi, the ΔSLO strain is twice less cytotoxic than the WT strain (25 and 33% versus 43 and 60 %, respectively) (Figure 3G). However, at both time-points, the ΔSLO strain still induced a significant cytotoxicity compared to the non-infected control (26 and 33 % versus 1.6 and 1.4 %, respectively), indicating that GAS produces cytotoxic molecules other than SLO. In conclusion, GAS yields dramatic cell damages within a few hours of infection of the decidua and SLO, together with other secreted factors, is involved in the toxicity.

**Figure 3.**
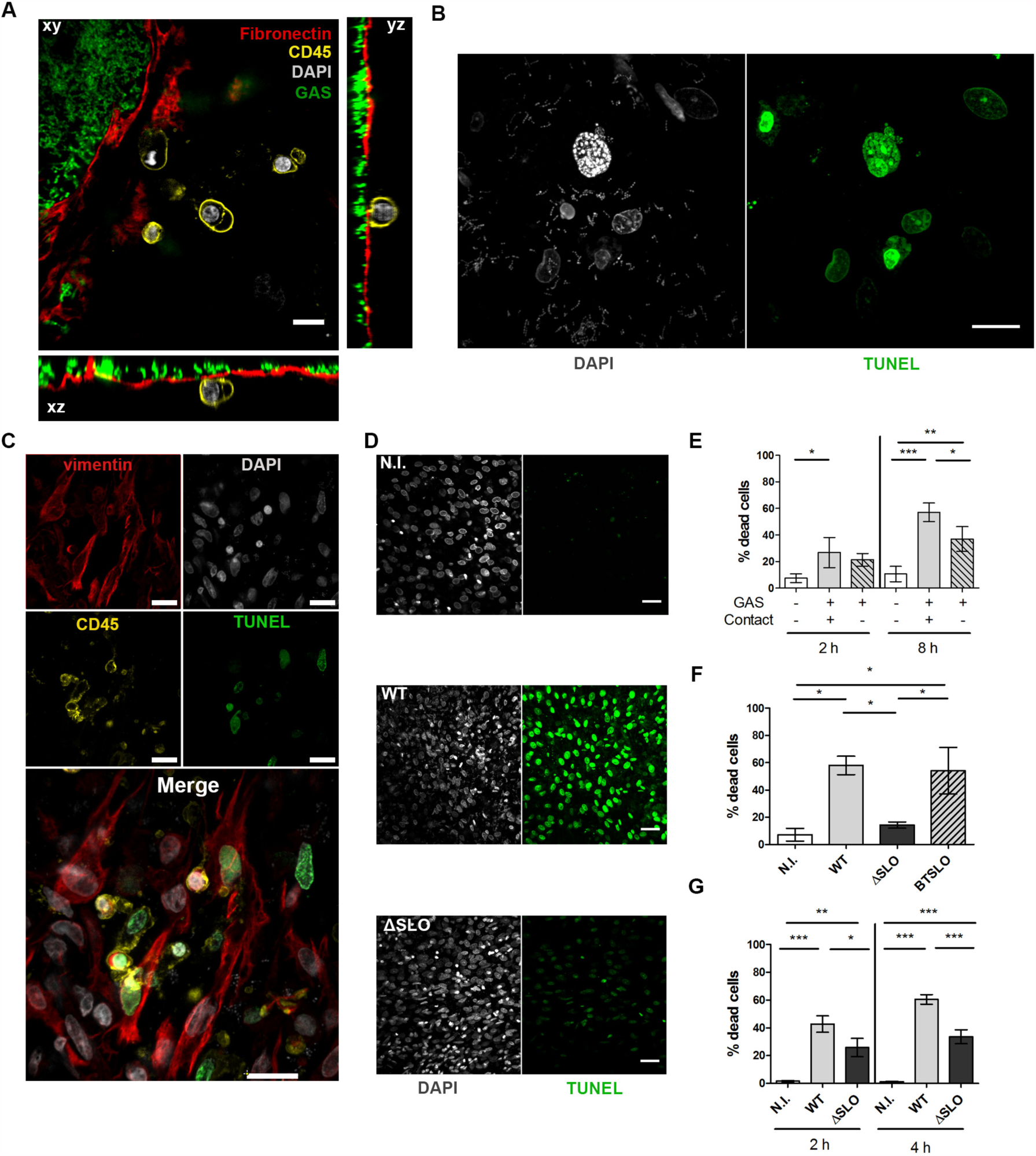
GAS induces stromal and immune cells death via processes involving SLO and other secreted factors. **A**, Orthogonal view of a tissue 5 h pi sc prestained with anti-fibronectin, red; anti-CD45 antibodies, yellow; WT-GFP bacteria, green; DAPI, grey. Scale bar: 10 µm, Magnification 40 X. **B**, Immunofluorescence of the decidua 16 h pi sc; DAPI, grey; TUNEL, green. Scale bar: 10 µm. Magnification: 63 X. **C**, Immunofluorescence of a tissue 16 h pi sc. Decidual stromal cells are vimentin^+^, CD45^−^ and immune cells CD45^+^; Vimentin, red; DAPI, grey; CD45, yellow; TUNEL, green. Scale bar: 10 µm. Magnification: 40 X. **D**, Immunofluorescence of tissue 4 h pi sc with the indicated strains; N.I., non-infected. DAPI, grey; TUNEL, green. Scale bar: 40 µm. Magnification: 20 X. **E-G**, Quantification of cytotoxicity by determination of the percentage of dead cells; **E**, on decidua with 3 subjects, in the absence, contact +, or the presence, contact -, of a Transwell® insert at indicated times; **F**, on primary decidual cells from three different subjects with indicated strains 4 h pi; N.I. non-infected; **G**, on decidua with 4 subjects with indicated strains, at indicated times. Statistical analysis **E** and **G**: Two-Way ANOVA with a Bonferroni post-test, *, p<0.05; **, p<0.005; ***, p<0.001; **F**: paired One-Way ANOVA, *, p<0.05.

### GAS limits the tissue immune response

The efficient bacterial multiplication questioned the host immune response. We first analyzed the expression of 133 unique immune genes related to “Inflammatory cytokines and receptors” and “Human innate & adaptive response” pathways of four independent subjects (Figure 4). Prior to infection, there is, as expected, low to moderate expression of the majority of the genes tested (Figure 4A, Table S1) (30). The 10% most highly expressed genes are related to “macrophage” and “cytokine-mediated signaling pathway”, reflecting the capacity of the decidua to generate an immune response. We next investigated the change in the host immune response at 8 h pi. Ten genes among the 133 tested showed a significant overexpression; they all encode cytokines and chemokines related to the ontology pathways, “acute inflammatory response” and “macrophages” (Figure 4A, Table S2). Among the 10 overexpressed genes, TNF, CXCL1, CXCL2, and IL23a are paradigmatic of an immediate early gene primary response (31,32). To assess the breadth of this immune response, we compared these data to a previously published response of human decidua to a GBS infection (33). At 8 h pi, in contrast to GAS, GBS induced the expression of 32 genes of these pathways, including 9 of the 10 genes overexpressed after a GAS infection (Table S2). Most genes specifically activated during the GBS infection are related to “cellular response to cytokine stimulus”. Noteworthy, in contrast to GAS, genes overexpressed during GBS infection include subsets of secondary response genes, such as NFKB1, CCR7, CD80, and ICAM1 (32). This comparison indicates that GAS impairs the tissue robust immune response that can be elicited by a *Streptococcus* pathogen which elicits milder endometritis (34). Strikingly, among the tested genes the only one overexpressed by a GAS and not by a GBS infection is IL10, encoding an anti-inflammatory cytokine. Altogether, these results show that GAS specifically induces a low and limited inflammation during the initiation of infection.

**Figure 4.**
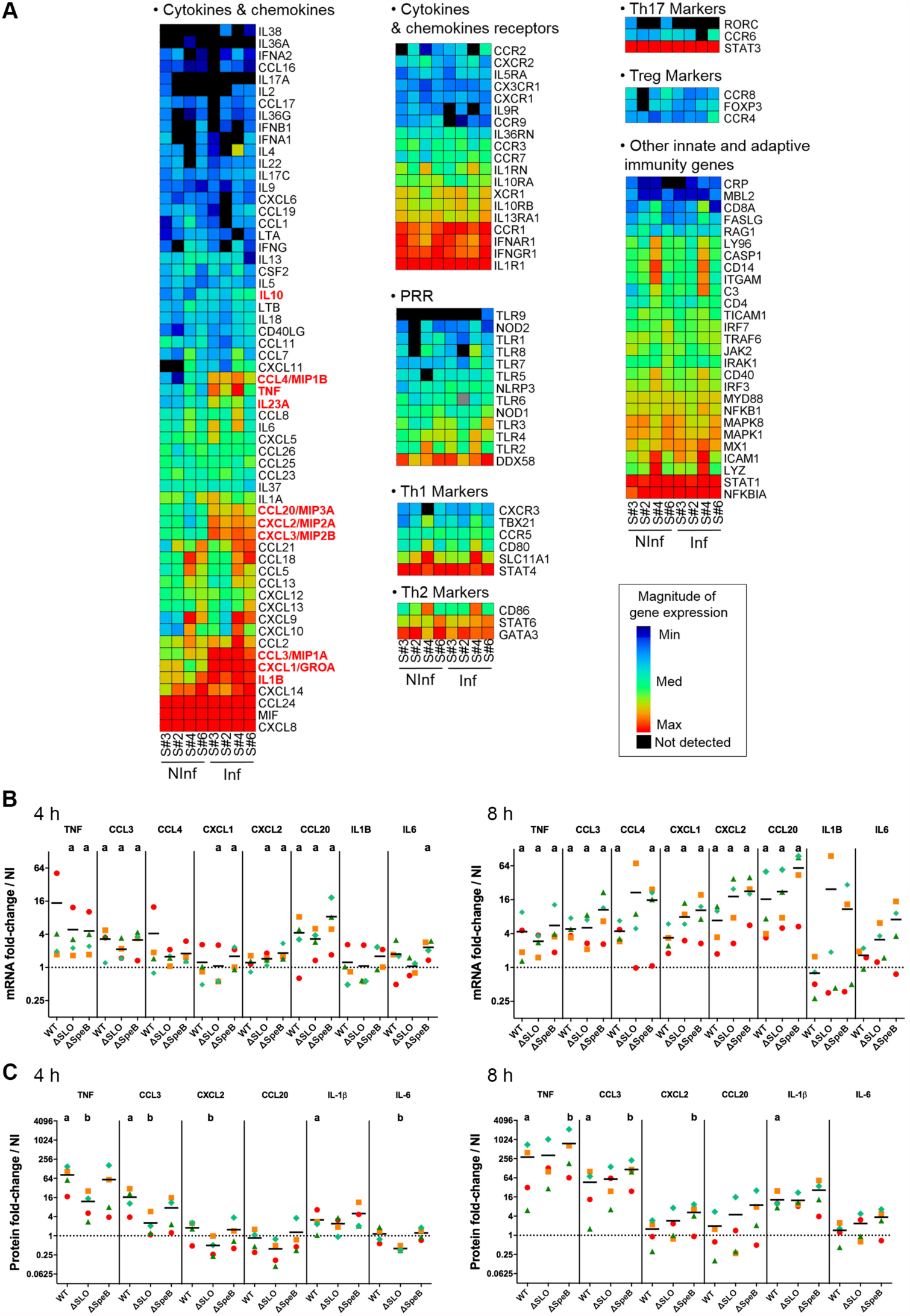
Host cytokinic responses to GAS infection. **A**, Heat-map of the results from the qRT-PCR screening using RT^2^qPCR “Inflammatory cytokines and receptors” and “Human innate & adaptive response” arrays on 4 subjects, non-infected (NInf) or infected 8 h sc with the WT strain (Inf). Cytokines significantly overexpressed in the infected compared to non-infected condition are shown in red. **B** and **C**, Fold-change variation of **B**, the expression of the immune-related genes after infection with the indicated WT or mutant strains for left, 4 h and right, 8 h pi compared to the non-infected expression. Statistical analysis: Mann-Whitney U test, one-way; a, p<0.05 as compared to non-infected. **C**, the concentration of the indicated cytokines compared to the non-infected condition at left, 4 h and right, 8 h pi on tissues from 4 different subjects. Statistical analysis: Friedman and post hoc pairwise comparison tests, a as in B b, p<0.05 as compared to WT. **B** and **C**, Axes are in log2. Symbols as in Figure 1I. Means are indicated with black lines. Dotted lines correspond to the value of 1.

To confirm the changes in gene expression, we quantified by qRT-PCR, on four additional tissues, the expression of five of the ten genes overexpressed after GAS infection at 4 h and 8 h pi, in uninfected and infected tissues (Figure S2A, 4B). In addition, since IL6 is the classical pro-inflammatory cytokine and although its expression did not appear modified in the arrays, we quantified its expression in the same conditions (Figure S2A, 4B). Overexpression of two genes, CCL3 and CCL20 occurs as soon as 4 h pi; at 8 h pi the expression of TNF, CCL4, CXCL1, and CXCL2 is also increased (Figure 4B). Notably, qRT-PCR assays confirm that IL6 is not significantly overexpressed after GAS infection, in contrast to the observed overexpression with GBS at 8 h pi (Table S2) (33).

Because GAS is also able to modulate *in vitro* the accumulation of cytokines (35–37), we looked if the overexpression of six cytokine genes translated in an increase in protein concentrations in tissue supernatants (Figure S2B, 4C). The basal level of accumulation of the six cytokines in the control tissues 4 h and 8 h pi shows the same pattern, independently of the subject, with high concentrations of IL-6 and CXCL2 (Figure S2B, Table S3,S4), known to be highly expressed in different subsets of decidual cells during pregnancy (38). After infection, the accumulation of TNF, CCL3 and IL-1β is significantly increased as soon as 4 h pi and even more at 8 h pi (Figure 4C, Table S3, S4). Interestingly, the overexpression of the CXCL2 gene at 4 h and of the CXCL2 and CCL20 genes at 8 h (Figure 4B) are not paralleled by an increase in the concentrations of these two proteins (Figure 4C), suggesting that GAS also controls the immune response at a post-transcriptional level.

GAS limiting the immune response at the transcriptional and post-transcriptional levels, we analyzed the potential roles of SLO and SpeB on that limitation. The activation of the tested genes was similar in the WT strain and the ΔSLO and ΔSpeB mutant strains (Figure 4B). In contrast, the SLO deficient strain generated a lower accumulation of TNF, CCL3, CXCL2 and IL-6, indicating that the presence of SLO increases these cytokines accumulation at 4 h pi (Figure 4C). Also, compared to the wild-type strain, the SpeB mutant had no effect on the accumulation of cytokines at 4 h pi, but induced a higher accumulation of TNF, CCL3, and CXCL2 at 8 h pi (Figure 4C). This indicates that SpeB tampers cytokine accumulation. Overall, GAS controls the immune response at the transcriptional and protein accumulation levels as soon as 4 h pi and SLO and SpeB are involved in the modulation of the immune response at the level of protein accumulation.

Since GAS influences the cytokine accumulation, we wondered whether it also modified AMP accumulation, and whether SLO or SpeB intervened. Infecting tissues with the WT-strain did not change the amount of AMP present in the supernatant (Figure S3, Table S5, Table S6), indicating that GAS infection does not elicit AMP accumulation. Infecting with the ΔSpeB strain yielded the same result as with the WT strain (Figure S3). In contrast, infecting with the ΔSLO strain induced significantly less human defensins 1 and 2 accumulation at 4 h pi. This suggests that SLO leads, as for cytokine accumulation, to increased human defensin 1 and 2 accumulation that is counterbalanced by other GAS-elicited signals.

## Discussion

GAS was, until the 20^th^ century, the major cause of infectious maternal death (39) while not being a colonizer of the vaginal flora (11) and puerperal fever is still responsible for > 75 000 maternal deaths per year, the most severe cases being caused by GAS (12). These data suggest that GAS has a particular propensity to elicit this disease. In addition, the onset of GAS puerperal fever occurs rapidly, the fever being established within two days following delivery. A comprehension of the crucial initial steps of GAS invasive infections and in particular puerperal fever is lacking. GAS possesses a large collection of virulence factors, but their role in the onset of these infections has not been characterized (7). Furthermore, the human specificity of GAS infections hinders the *in vivo* analysis of the initial steps. Using the original conjunction of *ex vivo* infection of human decidua, bacterial genetic manipulation, state-of-the-art image and immune response analyses, we demonstrated i) GAS adherence to and specific growth on the tissue within the first hours of infection, ii) GAS ability to invade the tissue by a proteolytic activity and maybe its persistence in immune cells and iii) GAS capacity to actively limit the host immune response (Figure 5). Our *ex vivo* setup, and particularly the live imaging, enabled us to dissect GAS capacity to establish decidua infection in real time and to approximate, with the presence of a flow, an *in vivo* situation.

**Figure 5.**
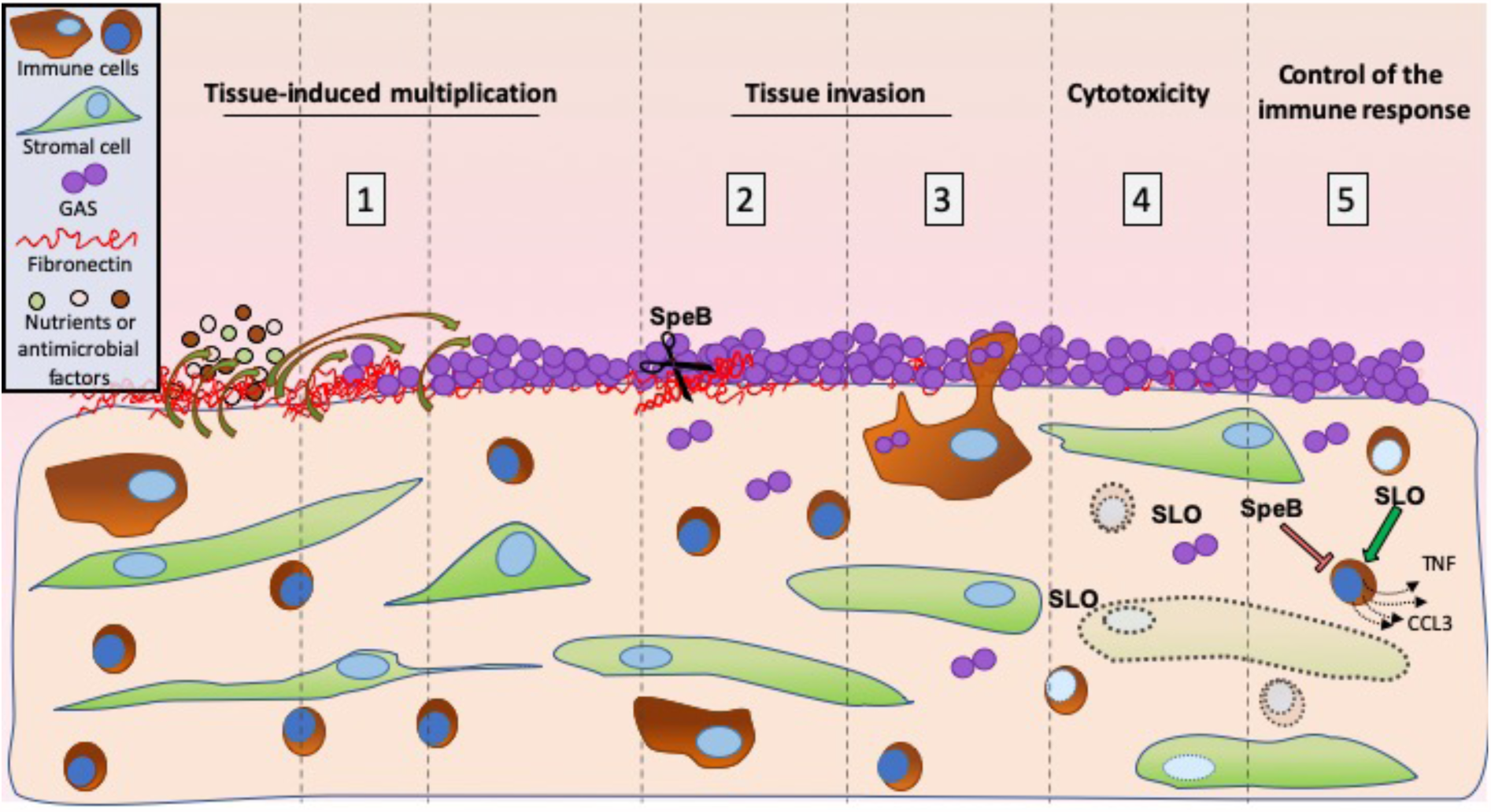
Schematic representation of the initial steps of decidua invasive infection by GAS. The successive steps are represented. 1) Decidua secretes anti-microbial peptides and nutrients (Figure 1I and S3) promoting GAS multiplication at the tissue surface (Figure 1C-H). 2) GAS invades the tissue in an SpeB-dependent manner, most likely through its capacity to degrade fibronectin (Figure 2A, B, C). 3) GAS is phagocytosed at the surface and is present inside decidual immune cells at least 6 µm below the surface and 4 h pi, suggesting GAS may use immune decidual cells to invade the decidua (Figure 2 D-E). 4) GAS induces the death of decidual stromal and immune cells in an SLO-dependent mechanism (Figure 3). 5) GAS limits the innate immune response, impairing the response amplification (Figure 4). Decidual immune and stromal cells are represented as observed in the tissue; bent arrows, nutriments, AMP, or growth signaling factors secretion and diffusion; dotted line objects, damaged cells; dotted arrows, cytokine secretion; red line, accumulation repression; green arrow, accumulation induction.

Our images demonstrate the presence of fibronectin and collagen at the surface of the decidua. It is unsurprising that GAS M28PF1 adheres to this tissue. Indeed, M28 strains, including M28PF1 (24), possess at least four fibronectin-binding proteins, Fba, SOF, SfbX and PrtF2, in addition to the six shared by strains from all genotypes. They also express one collagen-binding protein, Epf, in addition to the shared pilus and SpyAD (40). Finally, RD2, a 32 kb DNA region quasi specific to *emm28* strains modify, via encoded virulence factors and regulators, the virulence potential of *emm28* strains, contributing to their association with puerperal fever (41). The composition of the decidua surface and all these factors may explain the capacity of GAS to adhere to this tissue, a critical step in invasive puerperal fever.

To our knowledge, our study provides the first estimate of GAS doubling time in contact with a human tissue. Roughly 80 minutes is in the range of doubling-times described for other pathogens *in vivo* (42,43) and given the rapidity with which puerperal fever occurs it probably reflects the growth during the initial colonization step.

That GAS grows on the tissue of many subjects and in many uninfected conditioned supernatants, whereas it does not in the tissue culture medium, indicates that the decidua secretes nutrients and/or activation signals. In a controlled medium deprived of asparagine, SLO-induced endoplasmic reticulum stress leads to the secretion by epithelial cells of asparagine that promotes GAS growth (44). However, this mechanism required bacterial-cell contact and the medium we used for infection, RPMI, provides the 15 amino acids for which GAS is auxotroph (45). Since growth also occurred in uninfected conditioned supernatants, the secreted nutrient(s)/growth signal(s) are most likely not asparagine and defining them is the object of ongoing research. GAS grows on some tissues and conditioned supernatants and dies in others, suggesting that nutrient and/or bactericidal product secretion varies from subject to subject, both before and during infection. Whereas hBD1 and hBD2 concentrations were identical in all tissues, some differences may exist in that of LL-37. However, even if they existed, this would probably have little consequences on GAS killing since *in vivo* SpeB inactivates LL-37 at the bacterial surface representing a resistance mechanism (46). Interestingly, infection did not modify AMP concentration in the tissue supernatant, suggesting that the tissues display an intrinsic antimicrobial activity and that it may be controlled by GAS.

During invasive infections, despite its absence of motility, GAS readily disseminates within tissue and eventually to blood vessels, demonstrating GAS capacity to invade tissues. However, the crossing initiation, integrating the presence of extracellular matrix proteins has not, to our knowledge, been documented so far. Our data demonstrate that SpeB has an important role during the initial invasion step. Since SpeB has a broad proteolytic activity, including degrading extracellular matrix proteins, its role in tissue invasion is most likely linked to this capacity, degrading for instance the fibronectin which overlays the decidua (8). Yet, considering the pleiotropic effects of the absence of SpeB on the GAS surfactome and on GAS virulence mechanisms (8) our data cannot exclude that additional roles of SpeB are involved in the decreased invasion of the SpeB mutant strain.

Decidua parietalis is not covered by epithelial cells, the bacteria are therefore directly in contact, below the extracellular matrix proteins, with stromal and immune cells. Resident immune cells were followed by live imaging. Notably, neutrophils are normally scarce in the maternal-fetal membranes in the absence of infection (47). Phagocytic cells were present at the surface as soon as 15 min pi. Our data indicate that intra-tissular phagocytic cells readily capture surface bacteria in the first minutes of infection and that GAS is present at least 4 h pi in these cells. Although macrophages make an important contribution to the early control of GAS infection in intra-peritoneal murine model of infection (48), that GAS invades epithelial and immune cells and can survive or even escape from them has been documented in *in vitro* studies or during human biopsy analyses (49–51). Thus GAS ability to exploit the decidual inflammatory response by surviving inside phagocytic cells may be a virulence mechanism by favoring tissue persistence or/and invasion.

GAS induces the cell death of half the cells in the decidua, stromal and immune cells. The ΔSLO mutant strain is less cytotoxic than the WT strain, supporting *in vitro* obtained conclusions (52) and the blebs we observe could be related to SLO-induced blebbing documented on different cell types (53,54). Also, the ΔSLO mutant strain is still able to induce decidual cell death, suggesting the cytotoxic action of other GAS secreted products, such as the Streptolysin S (55). Hence, GAS cytotoxicity could account for phagocytic cells failure to cleanse the bacterial infection (56).

Dense macrophage and neutrophil infiltrations are observed during severe invasive GAS soft tissue infections in humans (50). A limitation of *ex vivo* infections is the absence of recruitment of immune cells (56). However, our setup allowed the analysis of the tissue intrinsic capacity to secrete cytokine and chemokines and consequently to recruit circulating immune cells. The expression of cytokines that recruit neutrophils, CXCL1, CXCL2, CXCL3, and macrophages, CCL3, CCL4, respectively (32,57), was increased within 4 to 8 h pi. Yet, after an LPS injection neutrophils infiltrate decidua via an IL1-dependent mechanism (47). However, after GAS infection in our setup the IL-1β mRNA regulation varied from subject to subject. The slight increase in IL-1β observed at the protein level for some subjects may result from an inflammasome activation (58). A single-nucleotide polymorphism in the enhancer-promoter region of IL1B gene has been associated with puerperal GAS sepsis susceptibility and could be responsible for the differences we observed (59).

No change was detected for chemokines receptors, other cytokines, or signaling mediators, especially from NF-κB family, indicating the absence of the response amplification. Furthermore, IL-10, an anti-inflammatory cytokine, was induced in response to GAS infection. This constrained inflammation may favor GAS infection initiation by delaying or decreasing the recruitment of immune cells and account for the GAS – puerperal fever association. The SLO and SpeB pro- and anti-inflammatory roles, respectively, are in accordance with previous conclusions that indicated a proinflammatory role for SLO (60) and that the genuine role of several GAS effectors is to impair the immune system, favoring GAS colonization (61). Interestingly, whereas *in vitro* SLO and SpeB optimal productions are not simultaneous, being during the exponential and the stationary phase of growth, respectively (62), the modified responses observed with the mutant strains indicate that they are sufficiently produced during the colonization and onset of invasion, as soon as 4 h pi, to fine-tune the immune response and to limit AMP accumulation.

Pathogens able to infect maternal-fetal interface are limited (63). *Listeria monocytogenes* causes foodborne disease with 12-20 times more prevalence in pregnant women, yet human decidua is a barrier to initial infection, inferring decidual stromal cells *per se* have defense mechanisms against *L. monocytogenes* (64). GBS is more prevalent in female reproductive tract than GAS but typically causes less severe maternal infection. In a model comparable to ours, where live GBS was in contact of decidua parietalis, GBS did not readily cross healthy term maternal-fetal membranes (65). GBS adherence to and penetration into the tissue was decreased at 8 h and 24 h pi compared to 4 h pi. In contrast to GAS, GBS, elicits robust increases of IL-6 and the chemokine CXCL8 (33). IL-6 has a pleiotropic effect on inflammation and CXCL-8 is one of the most important chemokine for the recruitment of neutrophils (57). Cell signaling molecules modulation was also noticeable only with GBS. Altogether, these data infer that GAS, in contrast to GBS, compromise the decidua innate response, including the upregulation of inflammatory mediators for neutrophil and macrophage recruitment.

In several monitored phenotypes, namely tissue invasion, cytotoxicity and immune response restriction, the virulence factors SLO and SpeB intervened. Other virulence factors which have been implicated *via in vitro* studies in non-immune cell entry, phagocytosis, inflammation modulation could be tested by our holistic approach (8,16). Transposon libraries could also be tested with the aim of finding novel colonization or invasion factors specific to this or other tissues. Our approach can also be applied to the study of the initial steps of other pathogen infections on human decidua, to compare with GAS efficiency, or on other human tissues such as the skin. Finally, GAS actively multiplies, hides as intracellular bacteria and diverts the host response during the very first hours following decidua infection. This rapidity may account both for the particular capacity of GAS, despite its *quasi-*absence from the female genital tract colonization, to elicit puerperal fevers and for their swift establishment. Our molecular decryption of GAS infection of the human decidua supports epidemiological conclusions for the necessity of maintaining and even increasing prophylactic measures (2,66) and the need to develop a vaccine to prevent life-threatening GAS infections such as puerperal fevers.

## Methods

### Strains and culture conditions

The strains used in this study are described in Table S7. The M28PF1 strain is a clinical isolate responsible for an endometritis (French National Reference Center (CNR) for Streptococci, https://cnr-strep.fr/) that was selected on phenotypic and genotypic bases from a collection of 50 *emm28* independent clinical isolates (24). GAS strains were grown under static condition at 37°C in Todd Hewitt broth supplemented with 0.2% Yeast Extract (THY) or on THY agar (THYA) plates. For GFP strains, the medium was supplemented with 10 µg/mL erythromycin.

### Genetic constructions and generation of GFP-producing and SLO- and SpeB-deleted mutant strains

The plasmids used in this study are described in table S7. The primers used for the generation of the plasmids and verifying the different strains are described in Table S8. The strains harboring an integrated inducible *gfp* gene were constructed as follows. We sequentially cloned into the pG1 vector (67), the Perm - *gfp* transcriptional fusion surrounded by the KpnI and HindIII sites from pATΩgfp (68) giving rise to pG1-Perm-gfp. The *lacA-lacR* intergenic sequence, enabling a single cross-over in M28PF1 and derivatives, was amplified using the primers F_lacA and R_lacA and cloned by In-fusion© into pG1-Perm-gfp digested with EcoRI, giving rise to pG1-lacA-Perm-gfp. The *erm* promoter was replaced by the tetO tetR Pxyl promoter region by amplifying it from pTCV_TetO (69) using the primers F_tetO and R_tetO and cloning it into pG1-lacA-Perm-gfp previously digested with EcoRI-EcoRV, giving rise to pG1-lacA-PTetO-gfp. The plasmid was checked by sequencing and transferred into M28PF1 or derivatives by electroporation as previously described (70). The correct localization of the construct was confirmed by sequencing the region using a primer outside the construct (R_extlacA) and one hybridizing to the vector, RP48. The uninduced GFP-producing strains, grew like M28PF1 in laboratory THY. The construct was stable even in the absence of erythromycin for at least twelve generations. The anhydrotetracycline (Sigma) concentration required for full expression of the GFP is 20-50 ng/mL and all bacteria are fluorescent after 90 minutes of induction, the fluorescence is stable for at least 6 h when the bacteria are in the stationary growth phase. GFP fluorescence is observable when GAS is intra-cellular. Induction of GFP does not affect strain growth.

The ΔSLO and ΔSpeB strains correspond to in-frame deletion mutants of the *slo* and *speB* genes and were obtained by homologous recombination of the plasmid pG1-SLO and pG1-SpeB following the same protocol as previously (14,71). The DNA fragments encompassing *speB* and *slo* were cloned in BamHI – EcoRI digested pG1 using the In Fusion cloning kit® (Clonetech). This led to deletion of nucleotides 72 to 1155 and 76 to 1690 for *speB* and *slo*, respectively. The back-to-the-wild-type (BTSLO and BTSpeB) strain was obtained during the second cross-over which leads to the mutant or back to the wild-type strain. The regions surrounding the construct were sequenced.

For all experiments, GAS strains were prepared as follows. Overnight cultures were diluted to an OD = 0.05 and grown in THY to the exponential phase (OD = 0.5), centrifuged and diluted in RPMI. For GFP expression, exponential phase bacteria were further diluted to an OD = 0.1 in THY supplemented with 10 µg/mL erythromycin and 20 ng/mL anhydrotetracyclin, grown for 1 h 30 min at 37°C and diluted in RPMI.

### Human tissue collection

Human placenta with attached maternal-fetal membranes were collected from healthy women with an uncomplicated singleton pregnancy, undergoing a planned cesarean delivery prior the onset of active labor at term (between 38 and 40 weeks of pregnancy). Except for per-operatively administered cephalosporin, women were excluded if prescription antibiotics were used during the two weeks preceding delivery. Table S1 indicates in which experiments each sample tissue was used.

### Maternal-fetal membranes processing

Within 15 min of collection, biological samples were processed in the laboratory. Maternal-fetal membranes were detached from placenta under sterile conditions, all similarly extensively rinsed in phosphate-buffered saline (PBS) and carefully examined. Pieces of membranes, at distance of placenta and of the remodeling zone overlying the cervix (13), free of surface blood clots, were cut and glued to Petri dishes or coverslips with a veterinary glue (Vetbond glue 3M, St Paul, MN), the fetal side sticking to the plastic or glass and covered with RPMI.

### In vitro *bacterial growth measurement*

RPMI was added to maternal-fetal tissues for 8 h at 37°C, 5% CO_2_; the supernatants (conditioned supernatants) were subsequently filtered and diluted 1/5 with RPMI. A thousand bacteria/mL was added to these diluted supernatants or to RPMI alone. After 8 h of incubation at 37°C, 5% CO_2_, solutions were serially diluted and plated. After 24 h at 37°C, CFU were counted.

### Primary decidual stromal cells (DSC) culture, infection and analyses procedures

Cells from three subjects were isolated from decidua parietalis, obtained from maternal-fetal membranes of nonlaboring women after a normal term pregnancy and cultured as previously described (14). One mL of GAS in RPMI at a bacterial density corresponding to a MOI of 100 was added to confluent DSCs for 4 h at 37°C, 5% CO_2_. Cells were extensively washed with PBS and fixed 15 min at 20°C with a 4% paraformaldehyde solution. Cytotoxicity was assessed using the cytotoxicity TUNEL staining following the manufacturer’s protocol (DeadEnd™ Fluorometric TUNEL System, Promega) and a LEICA DMI 6000 with a 20 µm objective (NA=0.5) for measurement.

### Antibodies

The following antibodies were used during this study: AlexaFluor 594 mouse anti-CD45 (Biolegend, clone M5E2), rabbit anti-Fibronectin (Sigma, #F3648), rabbit anti-vimentin (Abcam, ab92547), mouse anti-Type IV collagen (DSHB, M3F7), rabbit anti-whole GAS, gift from I. Julkunen. Secondary antibodies used were Alexafluor-594-coupled anti-rabbit and Alexafluor-647-coupled anti-mouse antibodies (Invitrogen).

### Experiments in static conditions (sc)

#### Immunofluorescence

8 cm^2^ or 1.5 cm^2^ of maternal-fetal membranes were glued to a 35 mm diameter Petri dish or on a 12 mm diameter glass coverslip respectively. When specified, tissues were pre-stained with anti-CD45 (1/100) for 30 min at 37°C, 5% CO_2_. Tissues were infected with 0.2 mL/cm^2^ of a solution of 1.7 10^8^ bacteria/mL in RPMI, supplemented by 10 µg/mL erythromycin and 50 ng/mL anhydrotetracycline (infection medium) when GFP strains were used. After infection, tissues were washed and fixed in formalin for 24 h, and then stored in 70% ethanol at 4°C until use. Fixed tissues were further cut into 0.5 cm^2^ pieces and glued to coverslips, permeabilized with 0.2% triton X100 (Sigma) for 30 minutes at 20°C. Blocking solution (PBS+BSA 3%) was added for 1 h at 20°C, and tissues were stained for 4 h at 20°C with blocking solution containing primary antibodies: anti-GAS 1/250, anti-fibronectin 1/250, anti-type IV collagen 1/50, anti-Vimentin 1/250. Tissues were washed with PBS and secondary antibodies and DAPI (1/1000) in blocking solution were added for 1 hour at 20°C.

Immuno-histo-fluorescence was performed on paraffin embedded tissues as described previously (18). 10 µm-thick slices were cut and stained with whole GAS rabbit antiserum (laboratory stock, 1/250) and with DAPI.

#### TUNEL assay

500 µl of RPMI were added to 1.5 cm^2^ tissue glued to a 12 mm glass coverslip. A hundred µL of M28PF1 at 5.10^8^ bacteria/mL was added directly to the tissue or in the upper chamber of a 6.5 mm Transwell® insert (polycarbonate, 0.4 µm membrane, Costar). For the non-infected condition, 100 µL of RPMI was added directly to the tissue. After the specified infection time, tissues were washed with PBS and fixed in formalin for 24 h, and then stored in 70% ethanol at 4°C until use.

Slight modifications were brought to the TUNEL manufacturer’s protocol. 0.5 cm^2^ pieces of tissue were fixed and glued to coverslips and permeabilized with 500 µL PBS with 0.2 % Triton X100 for 30 min at 20°C. A hundred µL of equilibration buffer (from the manufacturer’s kit) was added on the tissues in a wet chamber for 15 min, liquid was removed and 50 µL of labelling solution was added for 3-4 h at 37°C. Tissues were washed twice with saline-sodium citrate 2X buffer (from the manufacturer’s kit) and stained with DAPI (1/2000) for 15 min. For double vimentin and TUNEL staining, TUNEL staining was performed first.

#### Sc image acquisition

The maternal side of tissues was placed facing a 35 mm high glass bottom µ-dish (IBIDI) and imaged *en face* using a Yokogawa CSU-X1 Spinning Disk (Yokogawa, Tokyo, Japan) coupled with a DMI6000B Leica microscope (Leica microsystems Gmbh, Wetzlar, Germany). Acquisitions were made with MetaMorph 7 software. For quantification of bacterial invasion, images were acquired with a 40X objective (NA=1.25), 0.3 µm z-step and approximatively 90 slices per stack were acquired. For cytotoxicity quantification, images were acquired with a 20X objective (NA=0.7), 1 µm z-step and for tissues approximatively 70 slices per stack were acquired. For measurements of bacterial thickness and surface coverage, 3 to 12 fields per tissue were analyzed.

#### Sc image treatments and data analysis

All images were processed with ImageJ. For cytotoxicity measurements, TUNEL and DAPI positive nuclei of images were segmented and automatically counted. At least 5 fields per condition were analyzed for tissues, and 10 fields for DSCs. To measure bacterial invasion, another ImageJ macro was used to segment bacterial particles on z-max projection image of GFP signal. For each bacterial particle, the axial localization was determined and compared to fibronectin and type IV collagen localization. Any bacterial particle whose signal was below 1.2 µm to fibronectin or collagen signals was considered as an invasion event. Five to 8 fields (153 × 169 × 30 µm) per conditions were analyzed. A GAS chain was considered to be on average 4 cocci, corresponding to the mean size of invading particles. We estimated the amount of invading chains by considering bacterial chains surface and thickness (mean thickness, 4 µm, bacterial coverage of a field: 90 %).

Bacterial layer thickness: 3D surface heat map was generated by a custom R code based on the plot-ly library.

A 3D image representation of the tissue was performed using Imaris 7.4 (Bitplane AG).

### Experiments in flow conditions (fc)

8 cm^2^ maternal-fetal explant were glued to a 35 mm Petri dish and stained with Draq5 (1/2000) and anti-CD45 (1/100) in infection medium containing DAPI (1/30 000), fc infection medium, for 30 min at 37°C, 5% CO_2_. Tissues were extensively washed with fc infection medium and 3 mL of GFP-producing bacteria (1.10^8^ CFU/mL) in fc infection medium were added to the tissues in a 37°C microscopy chamber for 45 minutes until 20-30 % of the tissue surface was covered by bacteria. Supernatant was then discarded, tissue washed with infection medium to remove unattached bacteria and a peristaltic pump with circulating infection medium bubbled with 95% O_2_ and 5% CO_2_ was activated in an open circuit at 0.3 mL/min.

#### Fc image acquisition

Images of infected tissues were acquired *en face* with an upright DM6000FS Leica confocal microscope (Leica microsystems Gmbh, Wetzlar, Germany) coupled with a Yokogawa CSU-X1 Spinning Disk (Yokogawa, Tokyo, Japan), with a 25X objective (NA 0.95, Leica microsystems). Acquisitions were made with MetaMorph 7 software and images were taken every 30 min for 3 h with a z-step of 2 µm. For long-term acquisition, axial drift was manually compensated after 3 h and acquisition was resumed for another 3 h.

#### Fc image treatments and data analysis

All images were processed with ImageJ. 3D drifts of live images were corrected with custom routines. For surface area measurements of bacterial colonization, bacteria were segmented based on a threshold on the maximum intensity z projection images of the GFP signal at the different time points, with the same threshold for each time point. For local thickness measurement of the GFP signals corresponding to bacteria, whole field was subdivided in region of interest and a custom ImageJ macro measured the full width at half maximum (FWHM) of the z-axis GFP signal. The thickness value obtained by measuring the FWHM was then converted in µm by measuring the mean FWHM of 20 single bacteria, considered to be 1 µm thick.

#### Computer code availability

Computer codes used for this work will be transferred upon request.

#### RNA extraction

Sc-obtained tissues were washed once and stored in TriReagents (Sigma) at −80°C until used. Total RNA was isolated using a Qiagen RNeasy Kit (Qiagen, Valencia, CA) according to the manufacturer’s instructions. The purity and concentration of total RNA were evaluated using a Nanodrop spectrophotometer (Thermo Scientific, Waltham, MA), by measuring the ratio of absorbance at 260 nm and 280 nm.

#### RT-qPCR analysis

RNAs were treated with deoxyribonuclease (Invitrogen, Life Technologies, St Aubin, France) to remove any contaminating DNA. Four µg of total RNA were reverse transcribed using random primers and M-MLV Reverse Transcriptase (Invitrogen), according to the manufacturer’s instructions. Quantitative PCR was carried out on a Light Cycler 480, 96 well apparatus (Roche Diagnostics, Manheim, Germany), with 160 ng of cDNA as a template, using the amplification kit SensiFAST SYBR No-Rox kit (Bioline, London, UK), according to the manufacturer’s instructions. The RT^2^ Profiler “Inflammatory cytokines and receptors” and “Human innate and adaptive response” arrays PCR Arrays (Qiagen) were performed according to the manufacturer’s instructions. Gene expression was normalized using a panel of 5 housekeeping genes: ACTB, B2M, GAPDH, HPRT1, RPLP0. ΔCt ranging from 0 and below to 6.0 were considered as maximum magnitude of gene expression, from 6.1 to 9.9 as moderate and from 10 to 15 as low. Primers for additional RT-PCR analysis were chosen using PRIMER3 software, based on published sequences (available upon request). Primers were obtained from Eurogentec (Angers, France) and used at 10 nM in the PCR reaction. In this series of experiment, the set of internal controls included the geometric mean of three different reference genes: SDHA, PPIA, and GAPDH.

Genes with a nominal *p* value ≤ 0.05 were considered to be differentially expressed. Genes showing > 2-fold variation were further considered in the analysis. Heatmaps were created using the MeV Package. Enrichr was used for pathways enrichment analysis, restricted to Gene Ontology (GO) terms and ARCHS4 databases.

#### GBS infection of maternal-fetal membranes microarray subanalysis

Data were extracted using GEO2R from Park *et al*. (33). After conversion of the Gene annotation using DAVID gene ID Conversion tool, fold-changes and *p* values for the 133 immune-related genes we previously analyzed with the RT^2^Profiler Arrays were retrieved from these 8 samples.

#### Protein and antimicrobial peptide analysis

Sc-obtained supernatants of explants were stored at − 80°C until use. The levels of IL6, TNF, CCL20, CXCL2, CCL3, and IL-1 β were measured with Bio-Plex^®^ custom Assay (BIORAD) using a Bio-plex 200. AMP concentrations were analyzed in the same supernatants by ELISA with the following kits: LL37 (Hycult,biotech), human beta defensin 1 (R&D, Novus) and human beta defensin 2 (Elabscience). The concentrations are reported as pg/mL medium. The samples were quantified in duplicate according to the manufacturers’ instructions.

#### Statistics

Data were analyzed by Prism 6 software (GraphPad Software, Inc. San Diego, CA) or Xlstat version 2018.1 (Addinsoft, Paris, France). When indicated, we used Two-Way ANOVA. We used Student’s t-test or non-parametric tests for quantitative variables as indicated in the text and Pearson’s chi-square for qualitative variables, as appropriate. A *p*-value <0.05 was considered to be significant.

#### Study approval

The study of the human maternal-fetal membranes was approved by the local ethics committee (Comité de Protection des Personnes Ile de France III, n°Am5724-1-COL2991, 05/02/2013). All participants provided written informed consent prior to inclusion in the study at the Department of Obstetrics, Port Royal Maternity, Cochin University Hospital, Paris, France.

## Supporting information

Supp figures 1 to 3, supp Tables 1 to 8, legends for Supp video, Supp references

## Author contributions

A.W., T.G., C.L., C.M. and A.F. designed experiments, carried out experiments and performed data analysis. C.Pl., F.G. and C.Po. provided essential reagents and scientific advice. C.M. and A.F. supervised the project. A.W., C.M. and A.F. wrote the manuscript.

## Acknowledgements

We thank Louis Réot for the RT-PCR screening, Karine Bailly of the Cochin Cytometry and Immunology (CYBIO) for the Luminex cytokine quantification, S. Brochet and Q. Cece, from AF and CPo’s laboratory and T. Meylheuc from the Imaging Facility at the Microscopy and Imaging Platform (MIMA2, INRA, Jouy-en-Josas, France) for assistance with *in vitro* experiments, E. Donnadieu, S. Barrin, E. Peranzoni, V. Feuillet and A. Trautmann for advice in the imaging set-up, for helpful discussions, I. Julkunen for the gift of anti-GAS antibodies, the Imag’IC facility of the Cochin Institute, all the personnel of the CIC Mère-Enfant Cochin-Necker. This work was supported by Université Paris Descartes (AW contract n° KL2UD) the Département Hospitalo-Universitaire Risks in Pregnancy (PRIDE 2014, AF & CM), INSERM, CNRS, Université Paris Descartes.

## Notes

The authors have declared that no conflict of interest exists.

