## Supplementary material for "*Streptococcus pyogenes* infects human endometrium by limiting its immune response": Supp figures 1 to 3, supp Tables 1 to 8, legends for Supp video, Supp references

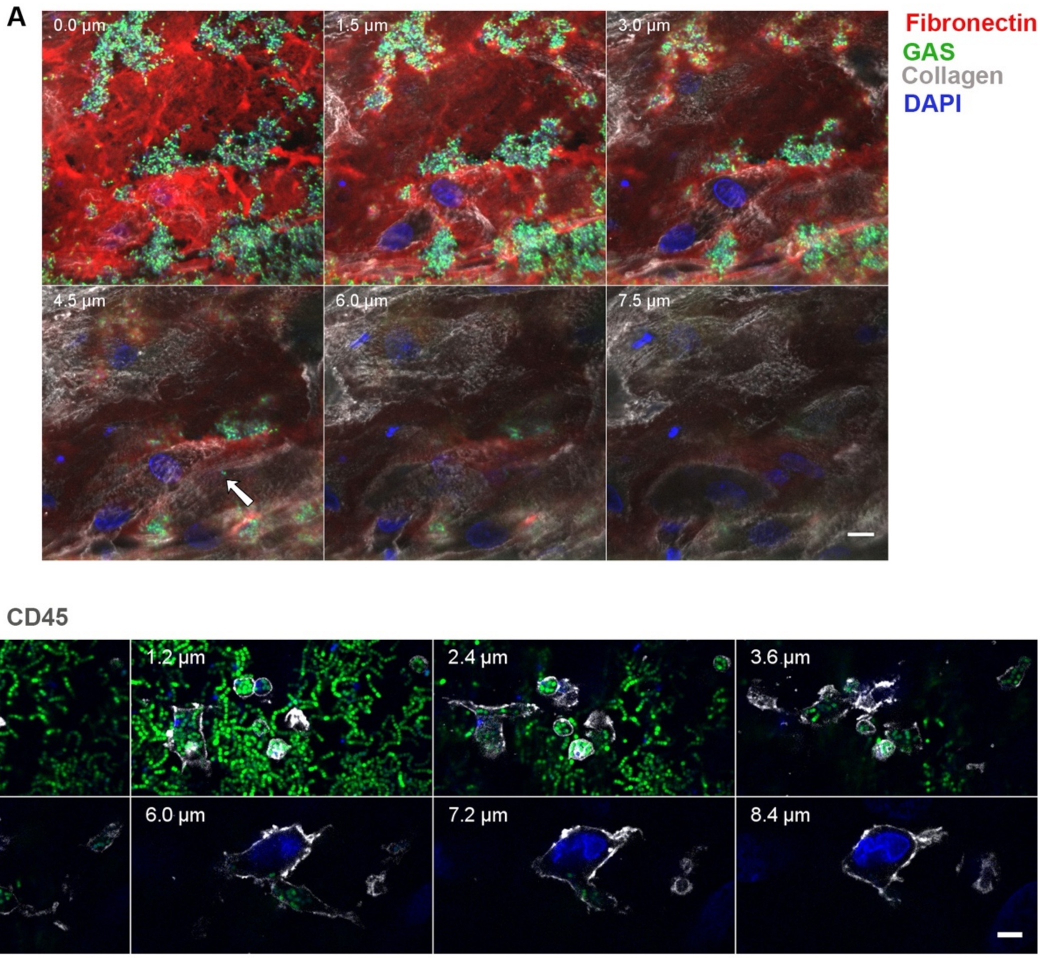


**Figure S1**. **GAS invades the tissue.** Montages of the same field at multiple depths. On both panels, the depth is indicated at the top left. 0 corresponds to the beginning of the GAS layer and is the first slice shown. **A**, ImageJ montage showing an example of GAS in the tissue. Tissue 16 h pi sc. Fibronectin, red; type IV collagen, grey; GFP-WT, green; DAPI, blue. A white arrow indicates the position of a GFP-WT coccus in the tissue. Scale bar: 10 µm. Magnification: 40 X. Same sample as in Fig. 1A and 3A. **B**, ImageJ montage of GFP-WT inside the tissue and within an immune cell. Tissue 3 h pi fc. Anti-CD45, grey; GFP-WT, green; DAPI, blue. Scale bar: 5 µm. Magnification: 100 X. Same sample as in Fig. 3E. All images are single slices acquired with a z-step of 0.3 µm.

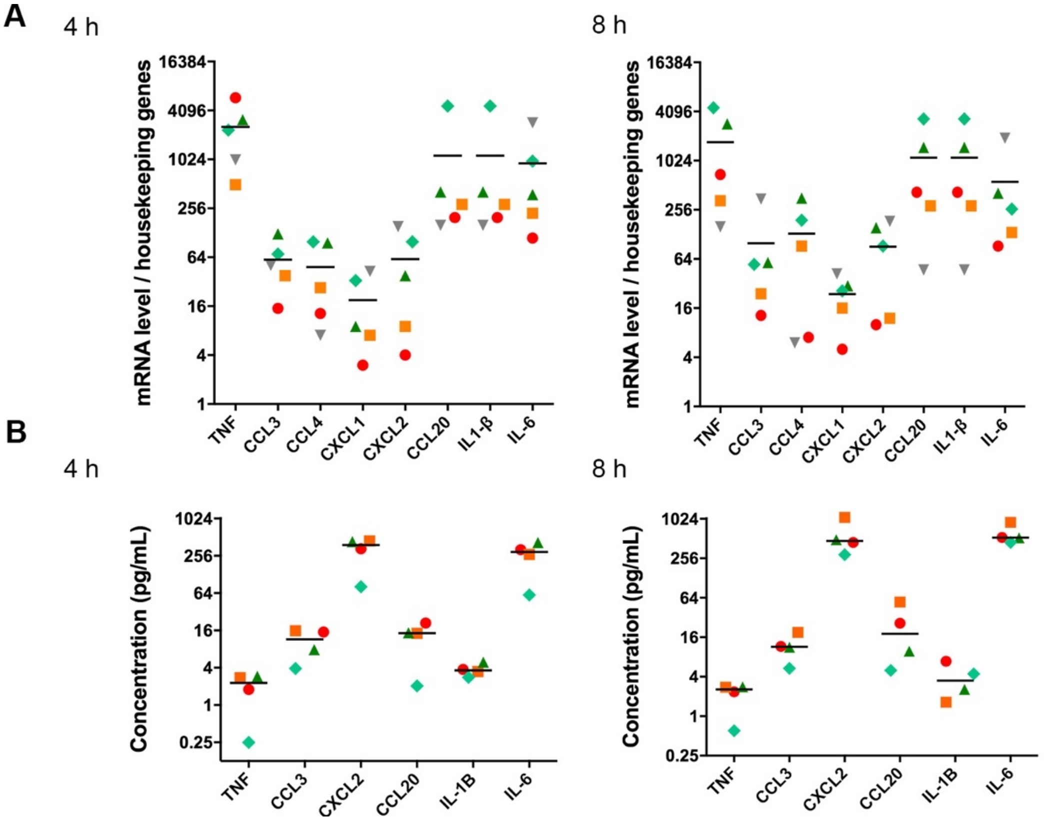


**Figure S2. Basal levels expression of cytokine genes (A) and accumulation of the cytokines in non-infected tissues (B). A**, Basal level of expression of the indicated immune-related genes in the non-infected conditions, at 4 (left panel) and 8 h (right panel) post-experiment starting point, compared to the housekeeping genes. **B.** Basal levels of accumulation of the indicated cytokine peptide 4 h (left panels) and 8 h (right panels) in the supernatant of non-infected decidual tissues. Symbols as in Figure 5B.

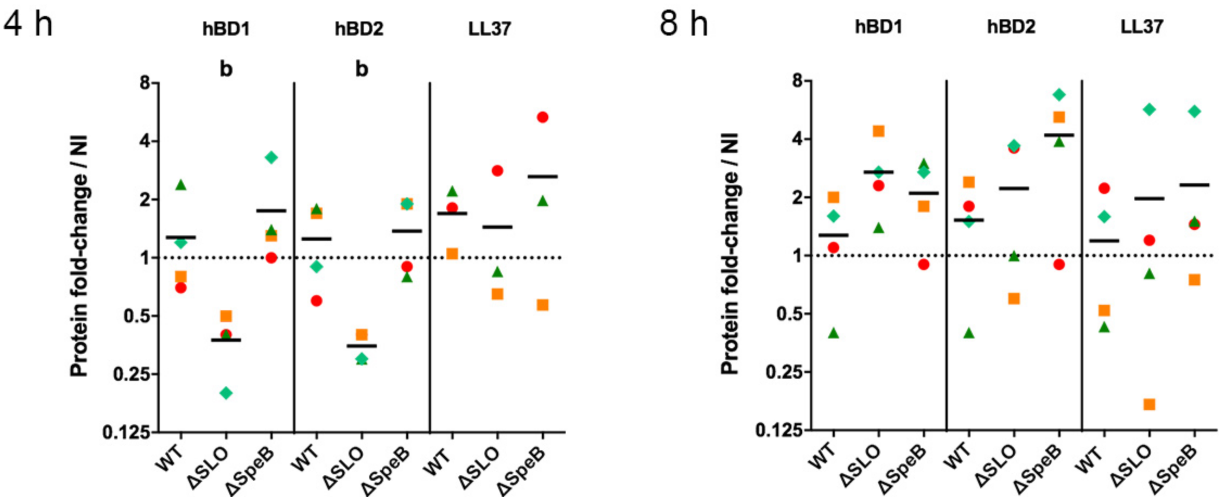


**Figure S3. Fold-change increase in the concentration of the indicated antimicrobial peptides compared to the non-infected condition (NI)**. Left, 4 h pi; right, at 8 h pi on tissues from 4 different subjects. Mean is indicated with a black line. Symbols as in Figure 1I. Statistical analysis: Friedman and post hoc pairwise comparison tests, b as in Figure 5C.

**Table S1.** List of experiments in which each tissue sample was used

| Subject[A] | Figure |
|---|---|
| S#2 | Fig. 1I, Fig. 3A-B, Fig. 4B, Fig. 5A |
| S#3 | Fig. 1I, Fig. 5A |
| S#4 | Fig. 1I, Fig. 5A |
| S#5 | Fig. 1C-D, Fig. 1F, Fig. 2C, Fig. 3E, Fig. 4E, Fig. S1B |
| S#6 | Fig. 1I, Fig. 5A |
| S#7 | Fig. 1H-J, Fig. 4C, Fig. 5B-C, Fig. S2A-B, Fig. S3, Table S3-6 |
| S#8[B] | Fig. 1I-J, Fig. 3B-C, Fig. 5B-C, Fig. S2A-B, Fig. S3, Table S3-6 |
| S#9 | Fig. 1C-D, Fig. 1G-J, Fig. 2A-B, Fig. 3B-D, Fig. 5B-C, Fig. S2A-B, Fig. S3, Table S3-6, Supp video 1 |
| S#10 | Fig. 1D, Fig. 1H-J, Fig. 5B-C, Fig. S2A-B, Fig. S3, Table S3-6 |
| S#11 | Fig. 5B, Fig. S2A |
| S#12 | Fig. 4G |
| S#13 | Fig. 4E |
| S#14 | Fig. 4E |
| S#15 | Fig. 4G |
| S#16 | Fig. 1A, Fig. 1H, Fig. 3A-C, Fig. S1A |
| S#17 | Fig. 1H, Fig. 4D, Fig. 4G |
| S#18 | Fig. S1D-F, Supp video 2 |
| S#19 | Fig. 1D, Fig. 1F, Fig. 4A, Supp video 3 |
| S#20 | Fig. 1B |
| S#21 | Fig. 4G |

[A]The numbering does not correspond to the order in which subjects were included in the study. [B] S#8 not included in Figures 1E and 1F because GAS did not grow in the presence of this tissue

**Table S2.** Comparison of the overexpression of genes involved in the immune response

| | GAS | | GBS | |
|---|---|---|---|---|
| | mRNA Fold-change | *p-value* | mRNA Fold-change | *p-value* |
| IL6 | 3.44 | 0.2443517 | 20.08 | 0.0000001 |
| **IL1B** | **6.06** | **0.0101306** | **7.79** | **0.0000004** |
| **TNF** | **40.37** | **0.0106377** | **32.25** | **0.0000016** |
| **CXCL2** | **29.34** | **0.0005470** | **10.46** | **0.0000044** |
| **CCL20** | **12.77** | **0.0005246** | **33.91** | **0.0000144** |
| CXCL8 | 1.90 | 0.0911197 | 4.80 | 0.0000319 |
| IL1RN | 0.94 | 0.9482826 | 3.96 | 0.0000747 |
| **CCL4** | **111.43** | **0.0032225** | **12.69** | **0.0000815** |
| IL1A | 1.83 | 0.4538192 | 6.85 | 0.0001290 |
| NFKB1 | 1.11 | 0.6764782 | 2.25 | 0.0001450 |
| ICAM1 | 2.08 | 0.4791674 | 5.01 | 0.0001730 |
| NFKBIA | 2.75 | 0.0500612 | 5.51 | 0.0001780 |
| **CXCL3** | **28.34** | **0.0000184** | **9.92** | **0.0001850** |
| TLR2 | 1.26 | 0.7816158 | 4.52 | 0.0002380 |
| TLR6 | 1.15 | 0.7529346 | 0.43 | 0.0017900 |
| CCL13 | 0.76 | 0.7698270 | 0.47 | 0.0018400 |
| NLRP3 | 2.36 | 0.0521823 | 4.39 | 0.0022200 |
| CCR7 | 2.62 | 0.1155378 | 3.36 | 0.0064500 |
| CCR1 | 2.00 | 0.0623347 | 0.36 | 0.0065000 |
| **CCL3** | **14.52** | **0.0009604** | **4.88** | **0.0072300** |
| CD80 | 1.61 | 0.5656690 | 3.08 | 0.0081700 |
| CD14 | 0.53 | 0.5807491 | 0.64 | 0.0084000 |
| IFNA2 | 0.84 | 0.8758766 | 0.71 | 0.0125000 |
| CCL2 | 2.58 | 0.2099790 | 2.38 | 0.0137000 |
| CCL8 | 1.71 | 0.5478146 | 3.10 | 0.0146000 |
| IL10RA | 0.56 | 0.2238125 | 1.55 | 0.0186000 |
| **CXCL1** | **14.52** | **0.0007640** | **3.20** | **0.0200000** |
| TICAM1 | 0.87 | 0.7283693 | 1.69 | 0.0226000 |
| **IL23A** | **9.47** | **0.0229023** | **13.52** | **0.0229000** |
| CXCL5 | 1.48 | 0.5061415 | 2.98 | 0.0233000 |
| IL37 | 0.64 | 0.1534900 | 0.68 | 0.0356000 |
| CD40LG | 2.38 | 0.3139426 | 0.74 | 0.0411000 |
| IL18 | 1.19 | 0.7368512 | 2.15 | 0.0510000 |
| LTA | 0.35 | 0.2617627 | 1.42 | 0.0511000 |
| TLR8 | 0.94 | 0.9689146 | 1.72 | 0.0609000 |
| CXCR2 | 0.51 | 0.3713690 | 0.54 | 0.0629000 |
| NOD1 | 0.78 | 0.5487833 | 0.60 | 0.0639000 |
| CXCR3 | 1.58 | 0.4743766 | 0.77 | 0.0803000 |
| IL9R | 0.23 | 0.1263553 | 1.27 | 0.0815000 |
| IRAK1 | 0.69 | 0.3244791 | 0.74 | 0.0861000 |
| TLR4 | 1.40 | 0.4818540 | 0.63 | 0.0879000 |
| TBX21 | 1.09 | 0.9344079 | 0.75 | 0.0966000 |
| RAG1 | 0.74 | 0.5642718 | 0.80 | 0.1000000 |
| CSF2 | 1.18 | 0.8418437 | 1.33 | 0.1030000 |
| IFNGR1 | 0.74 | 0.3574953 | 1.25 | 0.1120000 |
| STAT3 | 1.21 | 0.4820490 | 1.27 | 0.1330000 |
| CXCL10 | 3.57 | 0.4609962 | 5.21 | 0.1360000 |
| TLR1 | 0.93 | 0.9379901 | 0.70 | 0.1380000 |
| CASP1 | 0.76 | 0.7458079 | 1.42 | 0.1440000 |
| CD86 | 0.94 | 0.9553241 | 0.64 | 0.1730000 |
| **IL10** | **4.73** | **0.0490744** | **1.43** | **0.2190000** |

In green and bold the genes significantly upregulated after infection with GAS and GBS compared to tissue non infected; in orange the genes significantly only upregulated after infection with GBS (1). In red and bold genes only upregulated after GAS infection. Several genes only overexpressed by GBS, such as NFKB1, ICAM, NFKBIA, TLR2, NLRP3, CD80, are related to the ontogeny pathways "cellular response to cytokine stimulus" (GO:0071345, *p* value= $2.4\ 10^{-31}$). Hits were classified in the decreasing order of *p-value* for GBS, until no gene was significantly differentially regulated for GAS and GBS. n=4 for GAS and for GBS.

**Table S3** Concentration of inflammatory molecules in the supernatants at 4 h

| | CT | WT | ΔSLO | ΔSpeB |
|---|---|---|---|---|
| CXCL2 | **323** (81-444 ±84) | **544** (158-1091 ±222) | **167** (42-441 ±92) | **340** (132-629 ±104) |
| IL1b | **3.7** (2.8-4.9 ±0.44) | **11** (5.2-25 ±4.5) | **9.7** (2.6-18 ±3.3) | **18** (5.7-39 ±7.4) |
| IL-6 | **267** (60-417 ±75) | **346** (46-778 ±159) | **106** (21-145 ±29) | **279** (107-393 ±68) |
| CCL3 | **11** (3.9-16 ±2.9) | **186** (40-481 ±102) | **31** (8-92 ±20) | **82** (18-250 ±56) |
| CCL20 | **13** (2-21 ±4) | **9.6** (2.2-23 ±4.6) | **3.5** (1.6-7.1 ±1.3) | **8.2** (5.1-11 ±1.2) |
| TNF | **1.9** (0.25-21 ±2.9) | **130** (31-23 ±287) | **23** (3.7-7.1 ±70) | **58** (6.9-11 ±161) |

Concentration in pg/mL. Results are expressed as: Mean (minimum-maximum ± standard error). 4 subjects.

**Table S4** Concentration of inflammatory molecules in the supernatants at 8 h

| | CT | WT | ΔSLO | ΔSpeB |
|---|---|---|---|---|
| CXCL2 | **576** (291-1077 ±172) | **932** (153-2311 ±482) | **1115** (487-2115 ±351) | **3078** (410-7190 ±1453) |
| IL1b | **3.9** (1.6-6.9 ±1.2) | **43** (19-69 ±10) | **49** (15-97 ±18) | **75** (27-154 ±29) |
| IL-6 | **602** (448-904 ±102) | **951** (222-2203 ±432) | **1202** (495-2127 ±401) | **2274** (359-4283 ±853) |
| CCL3 | **12** (5.4-19 ±2.8) | **616** (18-1908 ±437) | **503** (73-761 ±158) | **1202** (279-2192 ±391) |
| CCL20 | **24** (5-55 ±11) | **33** (1.6-85 ±18) | **33** (3.1-79 ±17) | **147** (13-429 ±98) |
| TNF | **2.1** (0.6-55 ±2.8) | **401** (17-85 ±1090) | **320** (83-79 ±622) | **938** (152-429 ±1798) |

Concentration in pg/mL. Results are expressed as: Mean (minimum-maximum ± standard error). 4 subjects.

**Table S5** Concentration of antimicrobial peptides in the supernatants at 4 h

| | CT | WT | ΔSLO | ΔSpeB |
|---|---|---|---|---|
| hBD1 | **2212** (313-3369 ± 661) | **2807** (379-6579 ± 1324) | **934** (69-1378 ± 305) | **2923** (1045-3848 ± 636) |
| hBD2 | **4599** (2376-6405 ± 885) | **5356** (1862-8029 ± 1423) | **6288** (1862-11719 ± 2049) | **963** (301-1773 ± 304) |
| LL37 | **1.1** (0.36-2.1 ± 0.51) | **1.3** (0.61-2.2 ± 0.38) | **1.2** (0.31-2.7 ± 0.54) | **2.2** (0.71-5 ± 0.97) |

Concentration in pg/mL. Results are expressed as: Mean (minimum-maximum ± standard error). 4 subjects

**Table S6** Concentration of antimicrobial peptides in the supernatants at 8 h

| | CT | WT | ΔSLO | ΔSpeB |
|---|---|---|---|---|
| hBD1 | **4365** (2888-5561 ± 552) | **5072** (1677-7020 ± 1169) | **11102** (6402-12948 ± 1571) | **8998** (5193-13317 ± 2207) |
| hBD2 | **15146** (7426-29730 ± 5162) | **39634** (9835-89457 ± 39634) | **13077** (3527-32273 ± 13077) | **25009** (7706-52619 ± 10216 |
| LL37 | **1.6** (0.56-3.2 ± 0.62) | **1.8** (0.26-4.3 ± 0.89) | **1.7** (0.49-3.2 ± 0.68) | **2.3** (0.91-3.2 ± 0.49) |

Concentration in pg/mL. Results are expressed as: Mean (minimum-maximum ± standard error). 4 subjects

**Table S7** Strains and plasmids used in this study

| Strains or plasmid | Relevant properties | Source or reference |
|---|---|---|
| *Streptococcus pyogenes* | | |
| M28PF1 | Wild-type representative *emm28* clinical isolate | (2) |
| ΔSpeB | M28PF1 lacking the *speB* gene coding for SpeB | This study |
| ΔSLO | M28PF1 lacking the *slo* gene coding for SLO | This study |
| BTSLO | Back-to-the-wild-type, the reverting strain of the ΔSLO construction | This study |
| M28PF1-GFP | M28PF1 with the integrated pG1-lacA-PTetO-gfp | This study |
| ΔSpeB-GFP | ΔSpeB with the integrated pG1-lacA-PTetO-gfp | This study |
| Plasmids | | |
| pG+host5 | Erm; ColE1 replicon, thermosensitive derivative of pGK12; MCS pBluescript | (3) |
| pATΩgfp | pAT28 derivative containing the *gfp* gene | (4) |
| pTCV_TetO | Plasmid containing the tetO tetR Pxyl promoter, inducible with anhydrotetracycline | (5) |
| pG1-Perm-gfp | pG+host5 containing the *gfp* gene from pATΩgfp | This study |
| pG1-lacA-Perm-gfp | pG1-Perm-gfp with the lacA intergenic region to allow stable integration in GAS genome | This study |
| pG1-lacA-PTetO-gfp | pG1-lacA-Perm-gfp with the Erm promoter replaced by the tetO tetR Pxyl promoter of pTCV_TetO. Can be integrated in GAS genome for anhydrotetracycline inducible *gfp* expression | This study |

**Table S8**. Primers used in this study for cloning and checking cloning

| Primer Name | Sequence* |
|---|---|
| F_tetO | GTGGAATTGTGAGCGGATAAC |
| R_tetO | **GGTACC**TTTTCACTCGTTAAAAAGTTTTGAGAATATTTTATATTTTTGTTCATGT AATCACTCCTTCTTAATCTGTTAACGCTACGATCTAGCT |
| F_lacA | ACATGATTACGAATTTCAACGACTTCGTATTTACCTT |
| R_lacA | ACACTCTTAAGAATTCGCGGTCATATCTGAGATGTT |
| R_extlacA | CCACCATGGGTCCTGATA |
| RP48 | AGCGGATAACAATTTCACACAGGA |
| SLO-F1 | GACTCTAGA**GGATCC**GGTGCCAAAGGGTTTAGAA |
| SLO-R1 | CTCAGGGGGATAAGAGCTGCCGTTAGTAG |
| SLO-F2 | TCTTATCCCCTGAGCCCATATGGTTCGAT |
| SLO-R2 | CATGATTAC**GAATTC**GGGACAGTTGGGGTCAAATC |
| SpeB-F1 | GACTCTAGA**GGATCC**GAGCATCTACTAGCCACAATA |
| SpeB-R1 | GGGTTAGCAAGAACAAATCC |
| SpeB-F2 | TGTTCTTGCTAACCCTTCAACGGTTACCAAAGTGC |
| SpeB-R2 | CATGATTAC**GAATTC**ATTAGTAGGCGTTGATGACC |

* restriction enzyme sites are highlighted in bold and sequences used for the In-fusion© cloning are shown in red.

**Supplementary video:**

**Supp video 1. GAS multiplication at the tissue surface**
Live confocal microscopy of the human decidua infected by GAS under flow conditions, acquired *en face*. GAS is in green. Magnification: 25 X. Scale bar: 20 µm. Time step = 30 minutes. The images are the same as in Figure 1C and are a z-max intensity projection.

**Supp video 2. Isolated GAS multiplication at the tissue surface**
Live confocal microscopy GAS is in green of the human decidua infected by GAS under flow conditions, acquired *en face*. Magnification: 25 X. Scale bar: 5 µm. Time step = 30 minutes. The images are the same as in Figure 1E and are a z-max intensity projection..

**Supp video 3. Immune cell blebbing and death after infection**
Live confocal microscopy of tissue infected under flow conditions. Intact nucleus, red (Draq5); permeabilized nucleus (dead cell), blue; CD45 (immune cells), yellow. Magnification: 25 X. Scale bar: 10 µm. Time step = 30 minutes. The images are a z-max intensity projection.
